# SANTA FE OXA: Self-assembled oxaliplatin nanomicelle for enhanced cascade cancer chemotherapy via self-sensitized ferroptosis

**DOI:** 10.1101/2023.11.13.566938

**Authors:** Jianbin Shi, Wenjing Ma, Shunzhe Zheng, Fengli Xia, Xinying Liu, Ayumi Kikkawa, Kaho Tanaka, Ken-ichiro Kamei, Chutong Tian

## Abstract

The clinical utility of chemotherapy is often compromised by its limited efficacy and significant side effects. Addressing these concerns, we develop a self-assembled nanomicelle, namely SANTA FE OXA, which is composed of hyaluronic acid (HA) conjugated with ferrocene methanol (FC-OH), oxaliplatin prodrug (OXA(□)) and glycol-coupled linoleic acid (EG-LA). Targeted delivery is achieved as HA binds to the CD44 receptors that are overexpressed on tumor cells, facilitating drug uptake. Once internalized, hyaluronidase (HAase) catalyzes the digestions of the SANTA FE OXA, releasing FC and reducing OXA(□) into an active form. Active OXA induces DNA damage, while simultaneously promoting intracellular hydrogen peroxide levels via cascade reactions. In parallel, FC disrupts the redox balance within tumor cells, inducing ferroptosis. The synergistic combination of cascade chemotherapy and self-sensitized ferroptosis therapy has demonstrated remarkable anti-cancer efficacy in both *in vitro* and *in vivo* models. Moreover, this SANTA FE OXA significantly mitigates the systemic toxicity commonly associated with platinum-based chemotherapeutics. Our findings suggest a compelling advancement in nanomedicine for enhanced cascade cancer therapy.

## 1. Introduction

Chemotherapy remains a cornerstone in clinical treatment of numerous cancers,^[1–3]^ with platinum-based chemotherapeutic agents, such as cisplatin, oxaliplatin (OXA) and carboplatin, being particularly prevalent in the treatment of cervical, ovarian, head and neck, and non-small-cell lung cancers (NSCLC).^[4–7]^ These agents exhibit their remarkable efficacy to bind DNA within cancer cells, thereby inhibiting cell replication and inducing cellular death. OXA, the third-generation platinum compound, is notable for its unique spectrum of activity and mechanisms of cellular damage recognition. It has been reported to produce fewer DNA adducts than cisplatin at equivalent doses,^[8]^ yet its exhibits greater cytotoxicity. Nevertheless, the clinical application of OXA is limited by its tumor selectivity and systematic toxicity. Moreover, the efficacy of platinum drugs is often compromised by cellular ‘detoxification’ mechanisms, such as reaction with high concentration of intracellular glutathione (GSH) in tumor cells, leading to drug resistance.^[9]^

To circumvent these challenges, advanced nano platforms and platinum (IV) prodrugs have been developed to enable targeted tumor delivery and reduction-sensitive release, aiming to reduce systemic toxicity and counteract drug resistance mechanisms.^[10–12]^ Despite their potential, nanomedicines have yet to fully realize their therapeutic promise, primarily due to the difficulties in achieving targeted delivery and controlled drug release.^[13–15]^ Therefore, to enhance therapeutic efficacy necessitates the integration additional therapeutic approaches and the development of advanced nano platforms. Therefore, leveraging ferroptosis, an iron-dependent form of cell death, has shown promise in augmenting the anti-tumor effects by disrupting cellular redox balance.^[16–18]^

Ferroptosis, is characterized by the accumulation of lethal lipid peroxides (LPOs), driven by Fenton reaction, which utilize iron ion to catalyze the conversion of hydrogen peroxide (H_2_O_2_) into highly reactive hydroxyl radical (•OH). Ferrous ion (Fe^2+^), in particular, are more catalytically active than ferric ion (Fe^3+^), and essential cofactors for lipid peroxidases (Lipoxygenase (ALOX) and Cytochrome P450 oxidoreductase (POR)),^[19,20]^ thus playing a pivotal role in the ferroptotic process. Targeted delivery of Fe^2+^ to tumor sites is therefore a promising strategy for inducing ferroptosis.^[21–24]^ Ferrocene (FC), a organometallic compound, can release Fe^2+^ in the presence of H_2_O_2_, making it an ideal exogenous source for inducing ferroptosis. Previous studies have reported that various nano delivery systems combining FC with chemotherapeutic agents (such as doxorubicin ^[25]^ and cisplatin^[26]^) or ferroptosis inducers (RLS3^[27]^ and fluvastatin^[28]^) can significantly enhance anti-tumor effects. Additionally, the efficacy of the Fenton reaction *in vivo* is contingent upon endogenous H2O2.^[29]^

Excitingly, platinum drugs can activate nicotinamide adenine dinucleotide phosphate oxidase (NADPH oxidase, NOX) within cancer cells,^[30,31]^ resulting generation of superoxide (O^2−^), which are subsequently converted into H_2_O_2_ by superoxide dismutase (SOD).^[30,31]^ This property makes OXA-based chemotherapy an ideal partner for ferroptosis, probiding a self-sustaining sourse of H_2_O_2_ for the Fenton reaction. Therefore, the combination of OXA and FC holds significant potential for improving cancer treatment outcome.

Furthermore, the emergence of tumor-on-a-chip (ToC) platforms has offered a transformative approach to drug evaluation, addressing the limitation of conventional two-dimensional (2D) cell-culture models that often failed to recapitulate the complex dynamics of *in vivo* environments.^[32]^ These microfluidic-based ToC platforms simulate the physiological conditions of tumor microenvironments, including three-dimensional tumor structures and fluidic dynamics, provide more accurate prediction of drug responses. ToC platforms have demonstrated significant potential in improving the correlation between preclinical and clinical outcomes, particularly in the evaluation of anti-cancer nanomedicines.^[33]^ They offer a dynamic microenvironment that is more representative of patient responces, allowing assessment of drug efficacy and resistance in controlled and reproducible manner.

Here we report a Self-Assembled NAnomicelle for TArgeted cascade chemotherapy via self-sustained FErroptosis therapy with OXA, namely SANTA FE OXA (Scheme 1). This has HA-FC-OXA-LA conjugates as a main building block, consisted with hyaluronic acid (HA) nanocarriers grafted with FC-OH, OXA(□) and ethylene glycol-linoleic acid (EG-LA) to specifically target CD44-overexpressing tumor cells. The design of micelles illustrates intracellular drug release upon exposure to hyaluronidase (HAase), allowing for the synergistic action of OXAw(□) and FC-OH to induce DNA damage and promote ferroptosis. We evaluate the efficacy of the SANTA FE OXA using various models, such as 2D cell culture, 3D dynamic ToC platforms and *in vivo* animal models to provide a comprehensive assessment of its therapeutic potential.

**Scheme 1.**
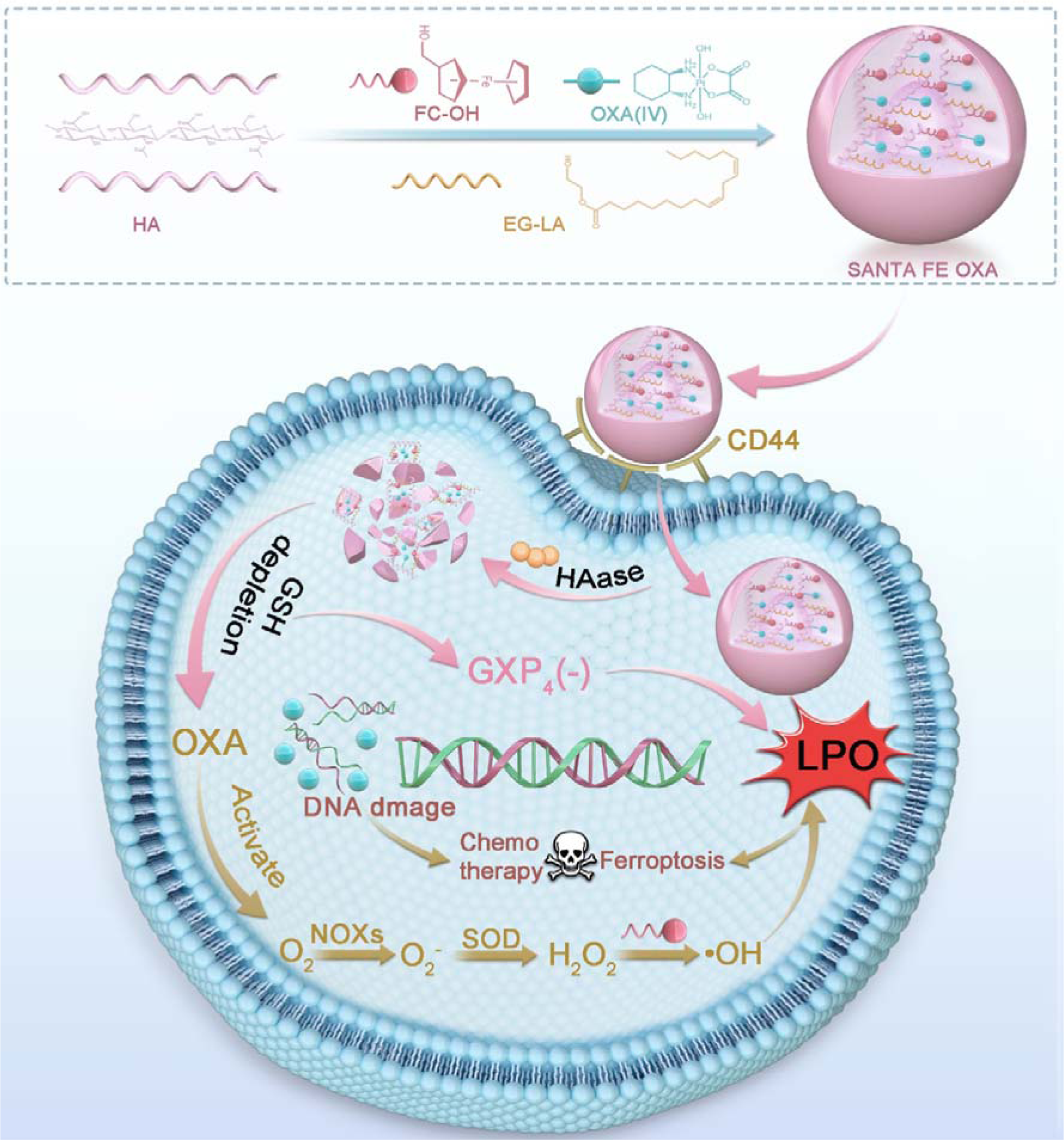
Self-assembled SANTA FE OXA and the ROS generation process for self-sustained ferroptosis and chemotherapy cancer therapy

## 2. Materials and Methods

### 2.1 Materials

Sodium hyaluronic acid MW 10 kDa, Fetal bovine serum (FBS), Dulbecco’s modified Eagle medium (DMEM), phosphate-buffered saline (PBS), trypsin, antibiotics, 3-(4,5-Dimethylthiazol-2-yl)-2,5-Diphenyltetrazolium Bromide (MTT), Hoechst 33342 and the others were purchased from Dalian Meilunbio. Oxaliplatin, Ferrocene methanol, Linoleic acid, *N*-(3-dimethylaminopropyl)-*N*′-ethyl-carbodiimide (EDCI), N-hydroxy succinimide (NHS), Deferoxamine mesylate (DFO), coumarin 6(C6), Vitamin C (VC) and the others were purchased from Shanghai bide pharm. Others Ki67 monoclonal antibody, GXP4 monoclonal antibody, β-Actin monoclonal antibody, Ki67 Enzyme-linked immunosorbent assay (ELISA) Kit, and the others were purchased from ABclonal Technology.

### 2.2 Synthesis of HA-FC-OXA-LA conjugates

OXA(□) was synthesized according to literature method,^[34]^ which was briefly described as (Figure S1, Supporting Information) suspension of oxaliplatin (100 mg, 0.25 mmol) in H_2_O_2_(30% (*w*/*w*), 5 mL) in the dark and stirred at 50 °C for 12 h. Pre-cooled methanol and ether were used to recrystallize and dry to obtain a white powder (86.8 mg)

For the synthesis of EG-LA. LA (3.5 mmol, 285.0 mg), DMAP (0.35 mmol, 43.1 mg) and EDCI (4.2 mmol, 386 mg) were dissolved in dry CH_2_Cl_2_ (10 mL) (Figure S2, Supporting Information). The mixture was stirred in an ice bath for 30 minutes before adding glycol and stirring at 25 °C for 24 h. Afterward, the reaction mixture was extracted 10% (*v*/*v*) citric acid and concentrated under reduced pressure. Silica gel chromatography (50:1 to 30:1 petroleum ether:ethyl acetate)

HA-FC-OXA-LA was synthesized using EDCI/NHS chemistry,^[35]^ and dissolved in distilled water (10 mL) with continuous stirring under an inert atmosphere of argon for 30 min at 25 °C. The HA-activated solution was further reacted with FC-OH (0.015 mmol 32.4mg) dissolved in DMSO (10 mL) at 80 °C for 18 h. The unconjugated free drugs were washed away by dialysis using a dialysis membrane (MWCO: 2000 Da) in excessive DMSO for 24 h and then excessive distilled water for another 24 h. The dialyzed products were lyophilized and stored at -20 °C for further use. Similarly, HA-FC-OXA was synthesized as described above. HA-FC activated solution was further reacted with OXA(IV) (0.015 mmol 64.4 mg) dissolved in DMSO (10 mL) for 18 h at 80 °C. The reaction product was dialyzed and lyophilized. Finally, HA-FC-OXA-LA was synthesized as described above. HA-FC-OXA activated solution was further reacted with EG-LA (0.0075 mmol 24.3mg). The reaction product was dialyzed and lyophilized.

### 2.3 Characterization of HA-FC-OXA-LA conjugate

The FC conjugation in the HA-FC conjugate was quantitatively determined using an ultraviolet-visible (UV–Vis) spectrophotometer. Absorbance values of FC (1–2 mg mL^-1^), HA (1 mg mL^-1^), and HA-FC (1 mg mL^-1^) were scanned at 300 – 650 nm.

Nuclear magnetic resonance spectrum (^1^H NMR) spectroscopy was used to characterize the HA-FC-OXA and HA-FC-OXA-LA conjugates, confirming the conjugation of OXA and EG-LA. The conjugates (5 mg mL^-1^) were dissolved in deuterated water or dimethyl sulfoxide-d6 for ^1^H NMR analysis using a 600-MHz Spectrometer. The resulting NMR spectra were analyzed with Mest Renova.

### 2.4 Preparation and characterization of the SANTA FE OXA

The critical micelle concentration (CMC) of conjugate micelles was analyzed using a published method.^[36]^

HA-FC-OXA-LA conjugate was dissolved in PBS to prepare SANTA FE OXA through its self-assembly properties. 12 mg of conjugates were dissolved in 2 mL PBS, then sonicated in an ice bath for 10 min (power: 100 W, working time: 2 s, intermittent time: 3 s), filtered with a 0.45-μm aqueous filter membrane, and the mic were stored at 4°C.

The morphology of the resultant Micelle was studied by a Tecnai G220 Transmission Electron Microscope. The hydrodynamic diameter, polydispersity index (PDI) and zeta potential of SANTA FE OXA was measured using Malvern Nano Zetasizer.

The static stability of SANTA FE OXA was evaluated by measuring the changes of particle size changes at 1, 2, 3, 4, 5, 6 and 7 days by the Malvern Nano Zetasizer. One mL micelle (6 mg mL^-1^) was mixed with 1 mL fetal bovine serum (FBS) and 8 ml PBS, and the solution was oscillated in a constant temperature shaker at 37 □. Samples were taken at 0, 0.5, 1, 2, 4, 6, 8, 12 h to evaluate the serum stability of SANTA FE OXA by measuring changes in particle size with the Malvern Nano Zetasizer.

In vitro release of OXA was determined using dialysis (*n* = 3). Briefly, SANTA FE OXA (400 μL) was immersed in a dialysis bag (MWCO: 2000 Da) containing 15 mL of dialysis medium PBS 7.4, vitamin C (VC) (10 mmol) and PBS 7.4, VC (10 mmol), HAase (100 units mL^-1^) and experiment was operated in a shaking bath at 100 rpm, 37 °C. At predetermined intervals, the external medium was collected and an equal volume of fresh dialysis medium was added. The OXA content in these samples was measured by HPLC at 220 nm using a mixture of 10% (*v*/*v*) CH_3_OH, and 0.1% (*v*/*v*) Formic acid in H_2_O.

### 2.5 In vitro GSH depletion

The consumption of GSH in vitro was monitored by the standard DTNB method. Briefly, The FC-OH (9.12 μmol, 19.7 μg), OXA (3.49 μmol, 15.0 μg), SANTA FE OXA (183.3 μg) and GSH (25 μmol) were individually mixed with 3 mL of PBS. The mixtures were labeled as follows: I: PBS, II:FC, III: OXA Sol, IV: FC+OXA Sol, V: SANTA FE OXA. These mixtures were then placed in a shaking bath at 100 rpm and 37°C. At predetermined time intervals, equal volumes) were added to the mixtures. After being kept in the shaking bath for 5 min at 100 rpm and 37 °C at 412 nm was measured by microplate reader (*n* = 3).

### 2.6 In vitro characterization of the Fenton reaction

The generation of •OH in vitro was monitored by the standard Crystal violet method. Briefly, the crystal violet solution (12 µM, 750 μL), H_2_O_2_ (100 mM, 500 μL) and FC methyl alcohol solution (FC 1 mM, 1 mL), OXA (411 μM, 1 mL), SANTA FE OXA (FC 1 mM, 1 mL) were mixed respectively, as follow: □: Crystal violet solution; □: Crystal violet solution + H_2_O_2_; □: Crystal violet solution + Methanol + H_2_O_2_; □: Crystal violet solution + OXA Sol + H_2_O_2_; □: Crystal violet solution + FC Sol + H_2_O_2_; □: Crystal violet solution + FC+OXA Sol + H_2_O_2_; □: Crystal violet solution + SANTA FE OXA + H_2_O_2_ The above mixture was kept for a shaking bath at 100 rpm and 37 °C for 2 h, and the absorbance at 580 nm was measured by microplate reader (*n* = 3).

Phenanthroline was adopted to verify the production of Fe^2+^. Briefly, SANTA FE OXA (FC 2 mM) was dispersed in 1 mM H_2_O_2_-containing acetic acid-sodium acetate buffer (0.2 M, pH 5.0). The mixture was incubated in a at 100 rpm and 37 °C for 2 h. Then, 900 µL of the reaction mixture was mixed with 100 µL of phenanthroline solution in an acetic acid-sodium acetate buffer, resulting in a final concentration of phenanthroline at 0.015% (*w*/*v*). UV absorption at 510 nm was measured by microplate reader (*n* = 3).

### 2.7 Cell culture

Human non-small cell lung cancer cells (A549), human triple-negative breast cancer cells (MDA-MB-231) and mouse Lewis lung cancer cells (LLC) were purchased from Dalian Meilunbio. The cells were cultured in DMEM medium supplemented with 10% (v/v) FBS and 1% (*v*/*v*) penicillin/streptomycin. The mouse normal fibroblast cell (NIH3T3) was purchased from Dalian Meilunbio. The cell was cultured in DMEM medium supplemented with 10% (*v*/*v*) FBS and 1% (*v*/*v*) penicillin/streptomycin. These cell cultures were maintained at 37 °C and 5% CO_2_ in a humidified cell incubator.

### 2.8 In vitro cell toxicity assay

The cytotoxicity of the SANTA FE OXA against A549, MD-MBA-231, LLC, and NIH3T3 cells was investigated in vitro using the MTT assay. Briefly, cells were seeded into 96-well plates at a density of 5000 cells/well and cultured for 12 h and the medium was discarded. The cells were incubated with different concentrations of the OXA Sol, FC+OXA Sol and SANTA FE OXA for 48 h. The cell viability was determined by the MTT assay. Cell viability was determined using the MTT assay. Cells without any treatment served as controls (*n* = 3). IC_50_ values were calculated and statistical analysis was performed using GraphPad Prism 8.

### 2.9 Western blotting

The level of GPX4 protein in treated LLC cells was assessed by western blotting. Simply, the LLC cells were seeded into 6-well plates at a density of 2 × 10^4^ cells/well and cultured for 12 h, which were treated with different formulations PBS, OXA Sol, FC + OXA Sol, SANTA FE OXA (OXA 50 μmol, FC 116 μmol). After 24-h incubation, the cells were collected and the protein was extracted with RIPA lysis buffer (Meilunbio). A BCA protein detection kit (Solarbio) was used to quantify total protein. The quantified proteins were then separated by SDS-PAGE and transferred onto polyvinylidene fluoride (PVDF) membranes. Subsequently, the membranes were blocked with blocking buffer containing 5% (*w*/*v*) skimmed milk and incubated overnight at 4 °C with primary antibody against GPX4 (1:1000). After incubation with the corresponding secondary antibody for 1 h, blots were visualized using enhanced chemiluminescence detection.

### 2.10 Design and fabrication of the microfluidic Chip

The chip has 3D structure channels. Each chip contains four channels (15 mm in length, 1.5 mm in width, and 0.5 mm in height), where the drug fluid will flow along the channel at a physiologically relevant velocity and pressure. The chip was fabricated using a published method.^[37,38]^

### 2.11 Construction of tumor-on-a-chip

A549 cells were collected and configured into a cell suspension at a density of 1 × 10^6^ cells mL^-1^. Then, 20 μL of the cell suspension was injected into each channel of the chip and cultured overnight at 37 °C with 5% CO_2_ for later use.

### 2.12 Cell uptake

Change the medium containing free C6 Sol, C6-Micelle or C6-Micelle + HA (10 mg mL^-1^), (C6 both at 200 ng mL^-1^) a flow rate of 10 μL min^-1^ or keep it static for 1 or 4 h. Stop the drug-containing medium and rinse the cells three times with PBS at 4°C. After adding 20 μL of 4% paraformaldehyde to each channel to fix them for 10 min at 25 °C, cells were rinsed three times with PBS at 4°C. Then, 20 μL of Hoechst33342 stain per channel was introduced, and cells were incubated for 15 min at 25 °C, and rinsed three times with PBS at 4 °C. 20 μL of anti-fluorescent quenching agent was added to each channel and finally capture images using a laser confocal scanning microscope. Process Images using ImageJ software (*n* = 3).

### 2.13 ROS assay

Cells were cultured dynamically with 10 μL min^-1^ medium containing PBS, FC Sol, OXA Sol, FC+OXA Sol, SANTA FE OXA, SANTA FE OXA + DFO, and SANTA FE OXA + VC for 4 hours (OXA 50 μmol, FC 116 μmol, VC 500 μmol, DFO 100 μmol). The cells were rinsed with PBS three times. Then the cells were incubated with DCFH-DA (10 μmol) for 20 min. After that, the cells were rinsed again with PBS three times. Finally, ROS production was observed using a laser confocal scanning microscope and analyzed using ImageJ software to process images (*n* = 3).

### 2.14 LPO assay

Cells were cultured dynamically with 10μL min^-1^ medium containing PBS, FC Sol, OXA Sol, FC+OXA Sol, SANTA FE OXA and SANTA FE OXA + DFO for 4 h (OXA 50 μmol, FC 116 μmol, DFO 100 μmol). The cells were rinsed with PBS three times. Then the cells were incubated with BODIPY (1 μmol) for 20 min. After that, the cells were rinsed again with PBS and LPO production was observed using a laser confocal scanning microscope and analyzed using ImageJ software to process images (*n* = 3).

### 2.15 Determination of GSH content

Cells were cultured dynamically with 10 μL min^-1^ medium containing PBS, OXA Sol, FC+OXA Sol, and SANTA FE OXA for 4 h (OXA 50 μmol, FC 116 μmol). The cells were rinsed three times with PBS. Then, the cells were collected by trypsinization and GSH and oxidized glutathione (GSSG) content was determined using glutathione kits (*n* = 3).

### 2.16 Evaluation of SANTA FE OXA with Tumor-on-a-Chip platform

A549 cells were collected and suspended at a density of 1 × 10^6^ cells mL^-1^. Then, 20 μL of the cell suspension was injected into each channel of the chip and cultured for 48 h at 37 °C with 5% CO_2_.The medium containing PBS, OXA Sol, FC+OXA Sol and SANTA FE OXA (OXA 50 μmol, FC 116 μmol) were dynamically perfused into the chip channel at and cultured at 37°C and 5% CO^2^ for 24, 48, and 72h. Ki67 monoclonal antibody and Ki67 ELISA Kit were used to investigate tumor cell proliferation on-chip.

### 2.17 In vivo antitumor efficacy in LLC subcutaneous xenograft mouse model

The LLC cells (1×10^6^ cells) were implanted subcutaneously in the flanks of C57BL/6 mice (body weight 20 g) respectively. The drug therapy was started on day 7 for LLC tumors. PBS, OXA Sol, FC+OXA Sol and SANTA FE OXA were given at a dose of 4mg/kg equivalent to OXA by tail injection every alternate day for 10 days (*n* = 5 for each group). The tumor volumes and body weights were monitored on daily basis. The animals were sacrificed when the average tumor size of the control 1200 mm^3^ in the control group. At the end of the treatment, the mice were sacrificed and the blood was collected and centrifuged at 4000 rpm for 10 min to obtain the serum for hepatic and renal liver function (alanine aminotransferase (ALT) and aspartate aminotransferase (AST)) and kidney function (creatinine (CRE) and blood urea nitrogen (BUN)) analysis. The tumor volume and burden were calculated as follows:

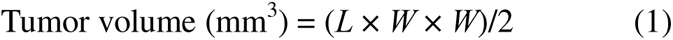

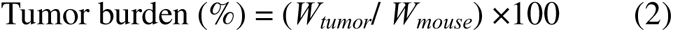

where *L* is the longest length of the tumor, *W* is the shortest width across the center of the tumor, *W_tumor_* is the weight of tumor, and *W_mouse_*is the weight of mouse.

### 2.18 Statistical Analysis

Statistical comparison between groups was analyzed with Student’s t test and one-way analysis of variance (ANOVA). Statistical significance was considered significant at *ns*: no significance, *\*P* < 0.005, *\*\*P* < 0.01, *\*\*\*P* < 0.001 and *\*\*\*\*p* < 0.0001. All results of the statistical analysis were expressed as mean ± sd.

## 3. Results and Discussion

### 3.1 Synthesis of HA-FC-OXA-LA conjugates

To provide the targeting delivery capabilities to SANTA FE OXA, HA with molecular weight of 10 kDa was chosen for its ability to conjugate multiple drugs, forming HA nanomicelle.^[39]^ HA is a natural acidic polysaccharide macromolecule, consists of two different saccharide units that furnish an abundance of functional carboxyl groups along its backbone, suitable for drug conjugation. HA has an inherent ability for CD44, which is often overexpressed in tumor cells, and is degraded by HAase present within the tumor microenvironments.^[40,41]^ Consequently, FC, OXA(IV) and EG-LA were conjugated to HA through EDCI/NHS-mediated reactions (Figure 1a). Initially, the carboxyl groups of HA was activated, allowing for a subsequently reaction with the hydroxyl group of FC-OH. In a similar way, HA-FC was activated, and reacted with OXA(IV) to obtain HA-FC-OXA. Finally, the EG-LA was reacted to obtain HA-FC-OXA-LA conjugates.

**Figure 1.**
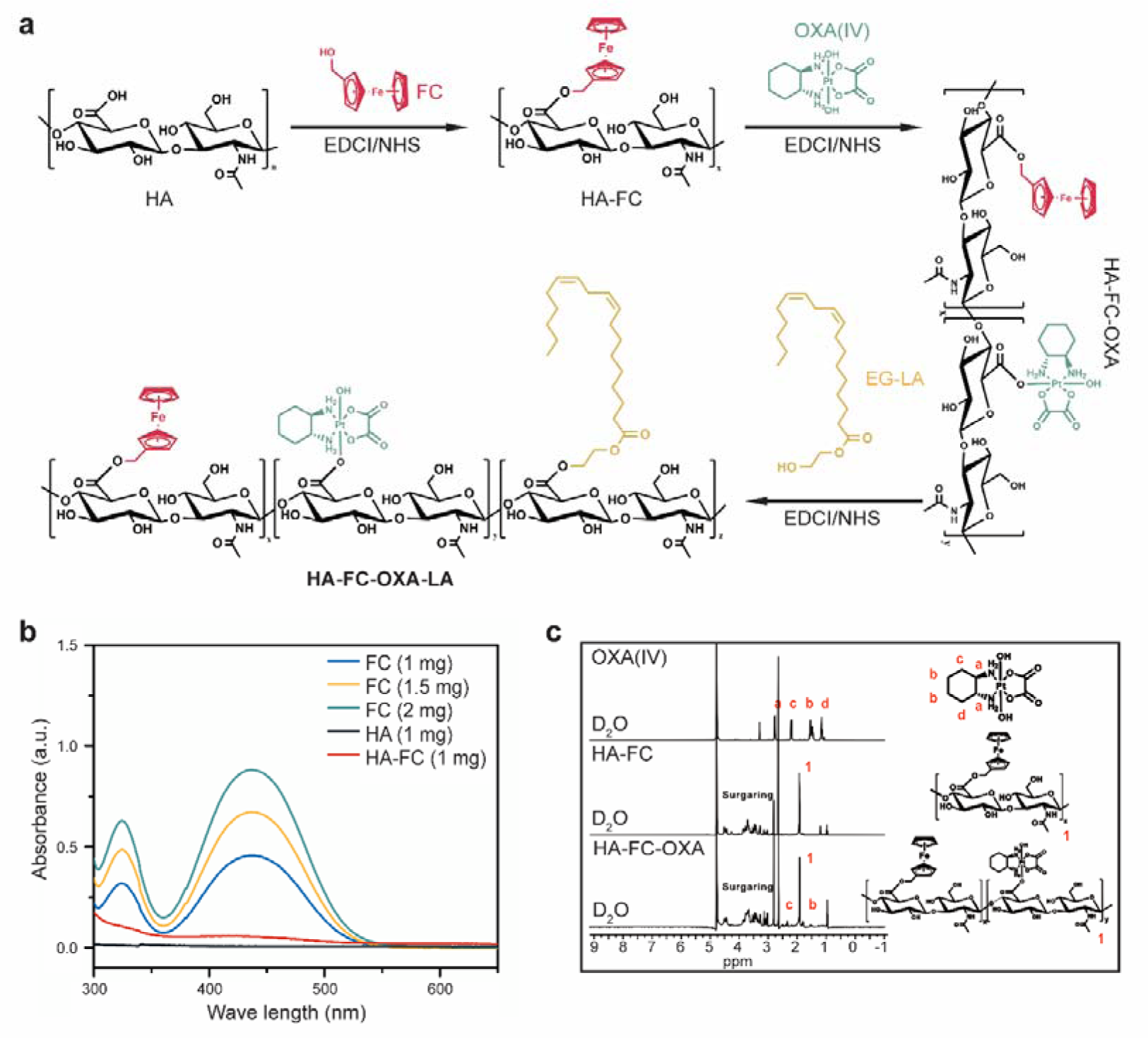
Synthesis and characterization HA-FC-OXA-LA conjugates. a) Reaction scheme depicting the conjugation of FC, OXA-□, and EG-LA in the HA backbone using EDCI/NHS chemistry sequentially. b) UV–Vis absorbance spectra of FC-OH, HA and HA-FC. Absorbance profiles are shown in the range of 300–650 nm wavelength. c) ^1^H NMR spectrum of HA-FC-OXA. Chemical shifts 1 (at 1.9 ppm) and b (at 1.5 ppm) represents N-acetyl group of HA and OXA, respectively.

The successful conjugation of these drugs to HA was confirmed by UV–Vis spectroscopy and ^1^H NMR spectroscopy, by comparison with standard FC-OH, OXA(IV) and HA spectrum (Figure 1b). The content of the FC in the HA-FC was found 22.3%, indicating successful ester bond formation between the FC-OH and HA. The conjugation of HA-FC with OXA(IV) and EG-LA were further analyzed and confirmed by ^1^H NMR spectroscopy. The ^1^H NMR spectrum of HA-FC-OXA displayed the major peaks of corresponding to OXA(IV) and EG-LA, in addition to the regular peaks of HA (Figure 1c, Figure S2, Supporting Information). The contents of OXA(IV) and EG-LA in HA-FC-OXA-LA were 8.25 and 4.00%, respectively.

### 3.2 Preparation and characterization of SANTA FE OXA

The self-assembly capacity of amphiphilic conjugates is a critical factor in SANTA FE OXA formation, with the critical aggregation concentration (CMC) serving as a key indicator.^[36]^ When the polymer concentration exceeds the CMC, it initiates self-assembly to form SANTA FE OXA with hydrophobic cores (Figure 2a). Following previously published protocol, the CMC of HA-FC-OXA-LA was determined at 7.76 × 10^2^ mg mL^-1^ (Figure 2b). Utilizing HA-FC-OXA-LA conjugate, SANTA FE OXA with two concentrations (6 and 12 mg mL^-1^) were prepared via an ultrasonic method. These amphiphilic HA derivatives spontaneously self-assembles into SANTA FE OXA in 1 × PBS. Both micelle formulations exhibited diameters ranging from 180.0 to 240.0 nm with PDI of 0.0886 – 0.144, as measured by Malvern Nano Zetasizer with conspicuous Tyndall effects (Figure S4, Supporting Information). Furthermore, their zeta potentials of the SANTA FE OXA are approximately -10 mV, reflecting the abundance carboxyl groups on their surface. Due to their similar properties, we selected the SANTA FE OXA formulation with a concentration of 6 mg mL^-1^ for subsequent analysis. Transmission electron microscopy (TEM) revealed a uniform spherical structure (Figure 2d), and the SANTA FE OXA exhibited stability in 1×PBS with or without serum (Figure 2e, f).

**Figure 2.**
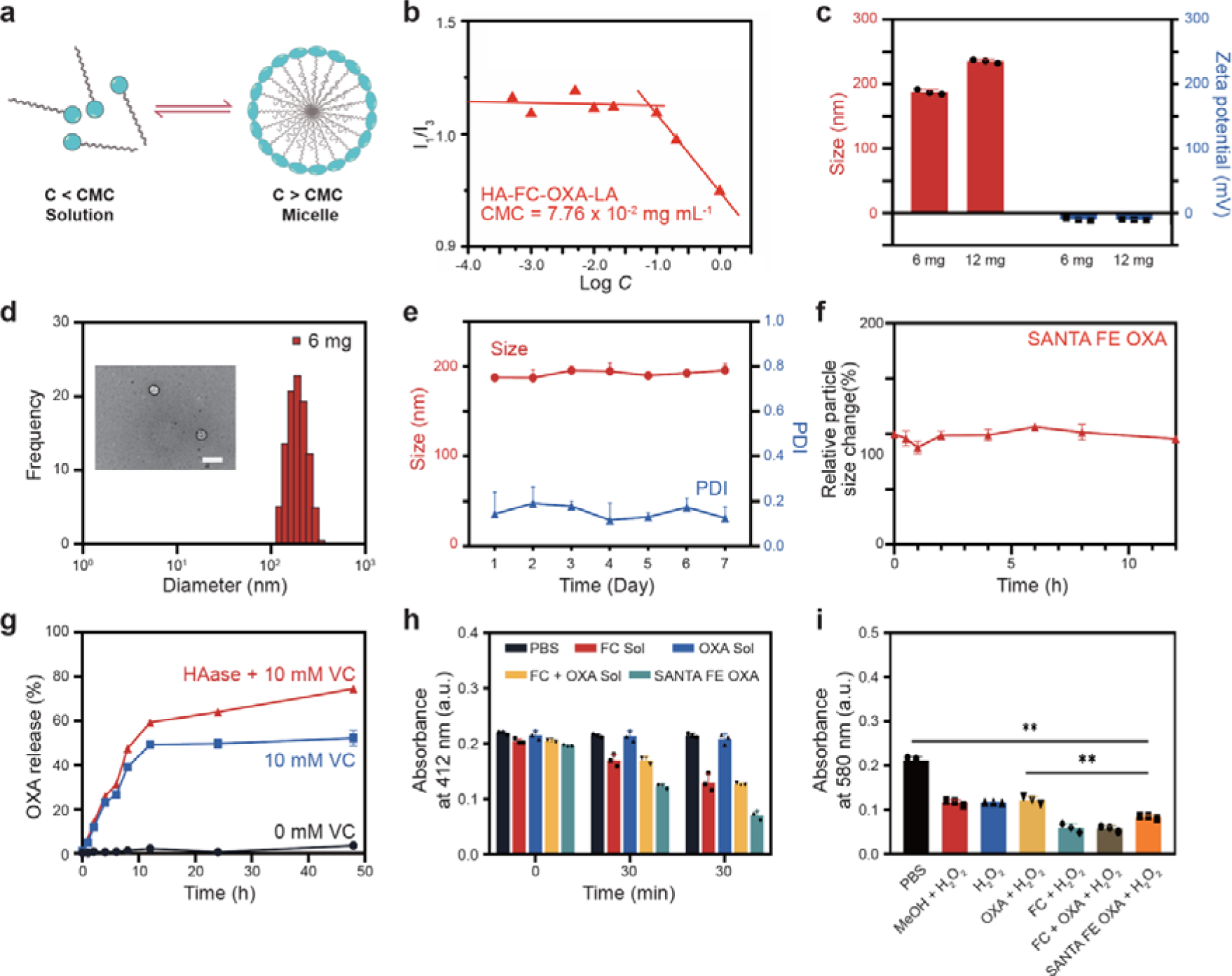
The preparation and characterizations of SANTA FE OXA. a) Schematic diagram of self-assembly principle of conjugates in water. b) Evaluation of critical aggregation concentration (CMC) of HA-FC-OXA-LA conjugates. c) The hydrodynamic diameter (Z-Average) and PDI of Micelles at different concentrations were measured by DLS. d) Transmission electron microscopic micrograph (Scale bar represents 500 nm) and intensity size distribution profiles of SANTA FE OXA (6mg mL^-1^). e, f) Evaluation of static (e) and colloidal stabilities (f) of SANTA FE OXA in PBS (pH 7.4) with or without 10% FBS (*n*=3). g) *In vitro* release profiles of OXA from SANTA FE OXA incubated in release media (PH=7.4) with 0 mM VC, 10 mM VC and 10 mM VC + 100 units HAase. h) GSH depletion induced by PBS, FC Sol, OXA Sol, FC+OXA Sol and SANTA FE OXA (*n* = 3). i) Hydroxyl radical (•OH) production induced by PBS, FC Sol, OXA Sol, FC+OXA Sol and SANTA FE OXA (*n* = 3). *ns*: no significance, *\*P* < 0.005, *\*\*P* < 0.01, *\*\*\*P* < 0.001 and *\*\*\*\*p* < 0.0001. Data are presented as the mean ± SD.

To investigate the reduction-responsive and enzyme-response drug release of SANTA FE OXA, we evaluated the release kinetics of OXA from the SANTA FE OXA at the presence of HAase and VC to simulate the intracellular environment of tumor cells (Figure 2g).^[42,43]^ In PBS with 10 mM VC, stimulating the intracellular reductive conditions, 52.2% of loaded OXA was released during 48 h. Furthermore, upon addition of HAase, simulating the intracellular enzymatic environments, 74.56% of loaded OXA was released during 48 h. In contrast, in the absence of HAase and VC, OXA is hardly released. The presence of 10 mM VC facilitated a gently controlled OXA release, which was accelerated by HAase due to the enzymatic degradation of the HA backbone. These results suggest that the SANTA FE OXA, leveraging the high levels of HAase and reductive glutathione (GSH) in tumor cells^[44,45]^, is primed for tumor-specific OXA release. This selective drug release is anticipated to be a minimal in normal tissues but enhanced in tumor tissues, potentially reducing the toxic side effects on healthy tissues, and improving the therapeutic index.

The GSH depletion capability of the SANTA FE OXA was explored by using 5,50-dithiobis-(2-nitrobenzoic acid) (DTNB) as an indicator, since the GSH depletion at tumor sites enhance the efficacy of ferroptosis therapies by increasing ROS accumulation and disrupting redox balance (Figure 2h, Figure S5, Supporting Information).^[46,47]^ Both FC and the SANTA FE OXA exhibited time-dependent depletion of GSH in contrast to the control. The micelle group showed more pronounced GSH depletion capacity, while Compared with the control group, OXA alone exhibited minimal GSH consumption and showed no significant change. The GSH depletion by FC is likely due to the presence of Fe^3+^,^[48]^ and the enhanced effect in the micelle group is attributed to the incorporation of OXA(IV), indicating that both FC and OXA(IV) have promising potential in reducing intracellular GSH levels and inhibiting GXP4 levels in tumor cells.

The SANTA FE OXA is specifically designed to activate the ferroptosis pathway by delivering FC and OXA. FC catalyzes the Fenton reaction, converting H_2_O_2_ into the highly reactive •OH, which initiate lipid peroxidation and ferroptosis.^[49]^ The crystal violet was used as indicator to verify the efficacy of this process.^[50]^ Therefore, we initially examined the •OH generation of various formulations (PBS, FC Sol, OXA Sol, FC+OXA Sol, SANTA FE OXA) through crystal violet degradation assays. In the presence of H_2_O_2_, neither OXA alone nor the control degraded the crystal violet solution; however, the addition of FC or the micelle resulted in a clear solution transition, indicating effective conversation of H_2_O_2_ into •OH (Figure 2i, Figure S6, Supporting Information). The relatively lower degradation rate of the micelle than free FC suggests a moderated reactivity of FC when conjugated to HA.

To quantify the release of Fe^2+^, we used the phenanthroline probe, which form a complex with Fe^2+^ to generate [Fe(phenanthroline)_3_]^2+^, detectable by its absorption band at 510 nm. It was evident that FC reacts with H_2_O_2_ to release Fe^2+^, corroborating the previous finding on the reactivity between FC and H_2_O_2_ (Figure S7, Supporting Information).^[51]^ The release of Fe^2+^ from the micelle, indicated by the formation of the [Fe(phenanthroline)_3_]^2+^ complex, signifies a rapid increase in •OH levels, further promoting ferroptosis.

Therefore, SANTA FE OXA exhibits a sophisticated dual-action mechanism for inducing tumor ferroptosis by releasing Fe^2+^ and depleting GSH, thereby disrupting the cellular tumor redox balance in a targeted manner. The selective activation of the SANTA FE OXA within the tumor microenvironments, due to the high concentration of H_2_O_2_ and GSH,^[44]^ enhances its anti-tumor efficacy while minimizing the risk of off-target side effects.

### 3.3 In vitro cytotoxicity evaluation

The cytotoxicity of various OXA formulations was assessed *in vitro* using the MTT colorimetric assay on tumor cells (A549, LLC, and MDA-MB-231). Across all cell lines, we observed a clear dose-dependent cytotoxic effects (Figure S8, Supporting Information). The IC_50_ of OXA Sol was lower than those of FC+OXA Sol, indicating a synergistic effect of FC and OXA in promoting ferroptosis and enhancing tumor cell death (Figure 3a, b, c). The SANTA FE OXA demonstrate a slightly reduced IC_50_ value compared with FC+OXA Sol, which can be attributed to three advantages: enhanced cellular uptake due to HA-targeting, increase intracellular OXA concentration, and the depletion of GSH by OXA(IV), which is crucial for maintaining cellular redox balance. Collectively, these advantages significantly boost the cytotoxic efficacy of OXA against tumor cells. In contrast, when assessing the cytotoxicity towards normal cells (NIH3T3), both OXA Sol and FC+OXA Sol demonstrated significant cytotoxicity against normal cells, whereas the SANTA FE OXA exhibited only minimal cytotoxicity, with an IC_50_ value 5.3 and 9.9 times higher than that of OXA Sol and FC+OXA Sol, respectively (Figure 3d). These results suggest that the SANTA FE OXA remains stable under normal physiological conditions, preventing premature drug release and thereby reducing toxicity to normal cells.

**Figure 3.**
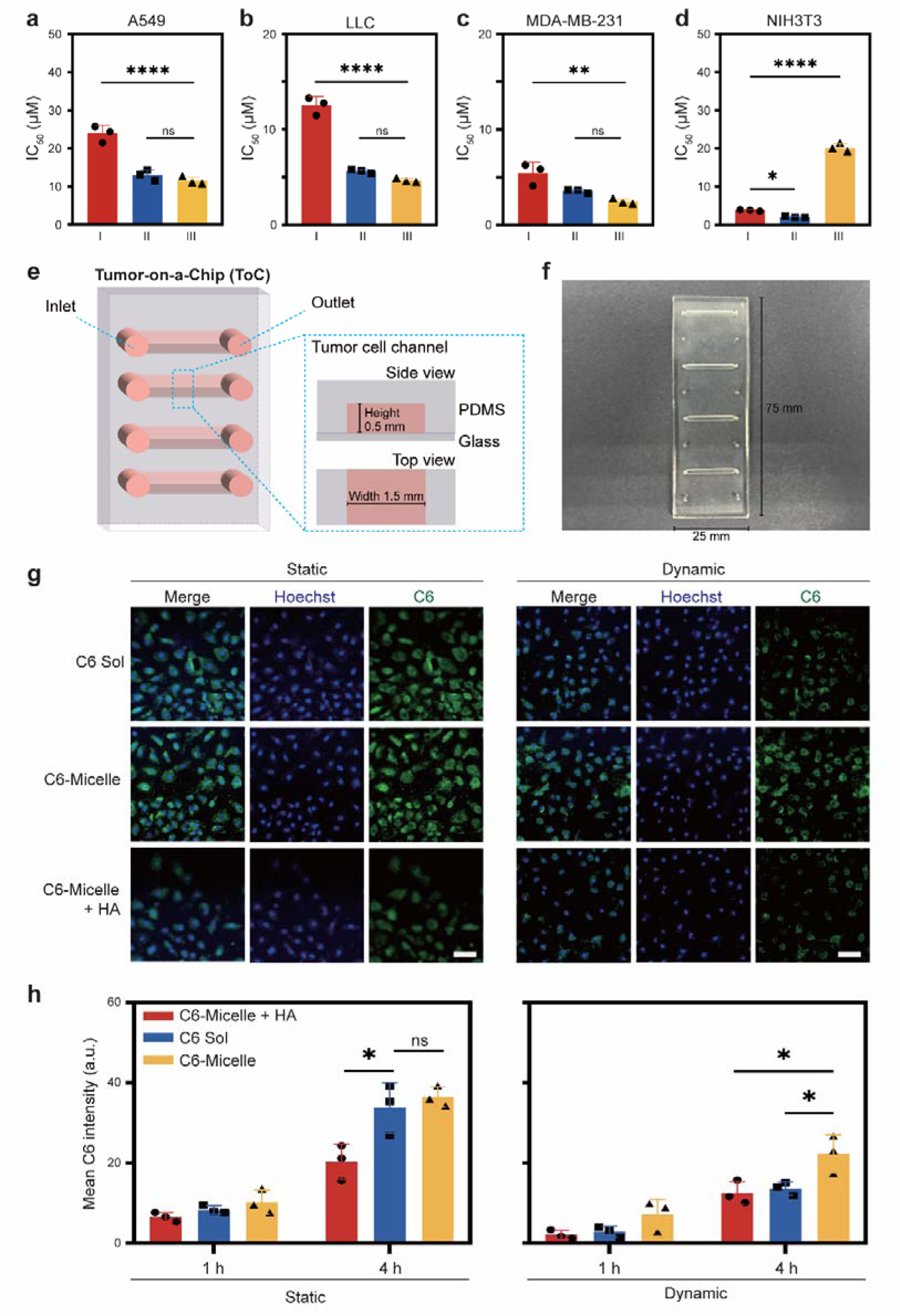
Evaluation of *in vitro* cytotoxicity and on-chip cellular uptake of SANTA FE OXA. a, b, c, d) Evaluation of the IC_50_ values of A549 (a), LLC (b), MDA-MB-231 (c), and NIH3T3 (d) cells treated with OXA Sol (I), FC+OXA Sol (II), and SANTA FE OXA (III). e, f) Illustration (e) and photograph (f) of a microfluidic Tumor-on-a-Chip (ToC) platform. g) Fluorescent micrographs of cellular uptake of C6 (Sol), C6-Micelles and C6-Micelles in the presence of HA (C6-Micelles+HA) in A549 cells cultured in a ToC platform under static/dynamic conditions at 4 h. Nucleus was visualized with Hoechst 33342 (10 µg mL^-1^). Scale bars represent 20 μm. h) Quantitative microscopic image analysis of cellular uptake of C6 (Sol), C6-Micelles and C6-Micelles+HA in A549 cells cultured in a ToC platform under static and dynamic conditions at 1 or 4 h. *ns*: no significance, *\*P* < 0.005, *\*\*P* < 0.01, *\*\*\*P* < 0.001 and *\*\*\*\*p* < 0.0001. Data are presented as the mean ± SD.

### 3.4 On-chip evaluation of cellular uptake

The efficacy of nano delivery systems is often contingent on the targeting cability and cellular uptake, which in turn is influenced by their physicochemical properties, such as size and surface charge^[52–55]^. To mimic *in vivo* delivery conditions, we employed a microfluidic dynamic ToC platform to assess the cellular uptake and intracellular drug release characteristics of SANTA FE OXA (Figure 3e, f, Figure S9, Supporting Information). The ToC platform, utilizing microfluidic chips, offers significant advantages over traditional 2D and even conventional 3D cell cultures in drug delivery research. It provides a more accurate and dynamic simulation of the complex tumor microenvironment, crucial for understanding drug behavior within the body. By studying multiple pathways for drug delivery under dynamic conditions, the ToC platform can provide insights into how factors such as flow rate, tumor microenvironment, and drug properties interact and affect drug delivery. We investigated SANTA FE OXA uptake under both static and dynamic conditions in a chip. Given the chip design and fluid dynamics during perfusion (with a viscosity approximately equal to 8.91 × 10^−3^ dyn s cm^−2^), we chose a perfusion flow rate of 10 µL min^−1^, resulting the shear force exerted of 2.38 × 10^−2^ dyn cm^−2^.

C6-labeled SANTA FE OXA (C6-Micelle) was prepared to visualize the cellular uptake of SANTA FE OXA. To confirm CD44-mediated endocytosis, cells were pre-treated with free HA to block CD44 (Figure 3g, Figure S10, Supporting Information).^[39]^ Under static conditions, C6-Micelles showed higher fluorescence intensity than C6 Sol at both 1 and 4 h (1.08-fold and 1.22-fold increase, respectively), indicating enhanced uptake via CD44-mediated targeting. The presence of free HA (C6-Micelle+HA) reduced the uptake of C6-Micelle in A549 cells, confirming the specificity of the HA-CD44 interaction. Under dynamic conditions, all groups showed decreased fluorescence intensity, suggesting that the shear stress reduced cellular uptake.^[56]^ This reduction is more pronounced in the C6 Sol group, which exhibited a 75.0% and 59.4% decrease at 1 and 4 h, respectively (Figure 3h). Furthermore, the binding affinity of the formulation to the cell membrane significantly influences the residence time parameter. Formulations with a higher binding affinity can resist the flow forces of the solution, thereby prolonging their interaction with cell membrane^[56]^. Consequently, the CD44-targeting ability of the SANTA FE OXA may offset the effects of flow rate, enhancing cellular uptake in the C6 Sol group.

### 3.5 Modulation of the Tumoral Redox Environment via SANTA FE OXA Formulations and In Vitro Drug Efficacy Assays

In this section, we aimed to identify the specific molecular pathways targeted by the SANTA FE OXA with tumor cells and elucidate their role in triggering ferroptosis-mediated cell death. To validate the synergetic induction capacity of SANTA FE OXA, we evaluated intracellular levels of ROS and LPO levels in A549 cells induced by dynamic administration of different formulations. This was conducted using the fluorescent ROS probe, 2’,7’-dichlorodihydrofluorescein diacetate (DCFH-DA) and LPO probe boron-dipyrromethene (BODIPY). Confocal laser scanning microscopy (CLSM) observations showed minimal fluorescence in cells treated with PBS or OXA, indicating relatively low ROS levels in these groups (Figure 4a, c, Figure S11, Supporting Information). In contrast, cells treated with FC exhibited a modest increase in fluorescence, reflective of limited endogenous H_2_O_2_ available for the Fenton reaction and subsequent production of •OH.

**Figure 4.**
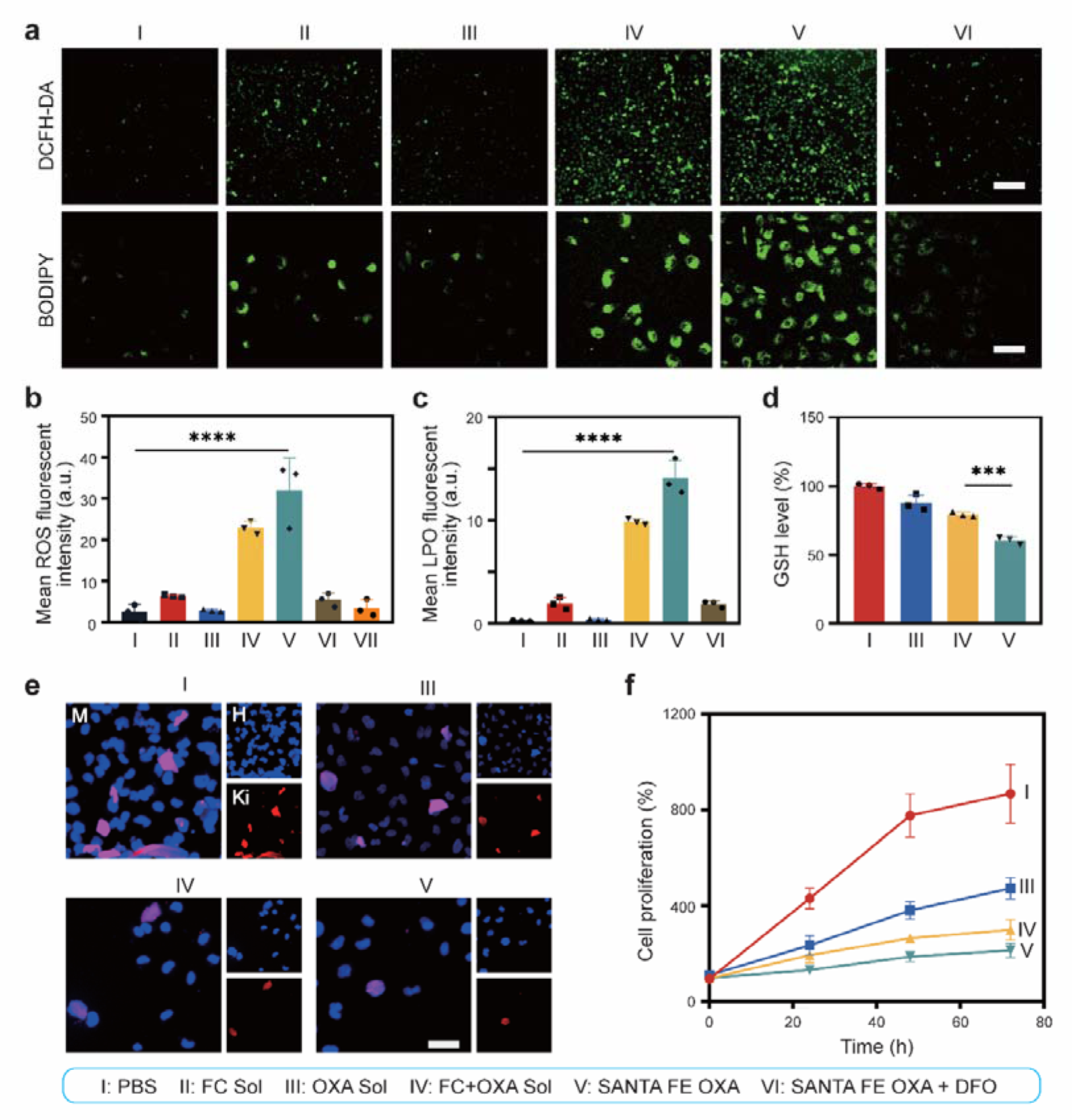
Evaluation of the Tumoral Redox Milieu via SANTA FE OXA Formulations by ToC drug efficacy assays. a) Intracellular levels of ROS and LPOs in A549 cells after dynamic treatments, visualized by DCFH-DA and BODIPY, respectively. Scale bars represent 100 μm. b) Quantitative microscopic image analysis of intracellular ROS levels in A549 cells (□, SANTA FE OXA + VC (Figure S11)). c) Quantitative microscopic image analysis of intracellular LPO levels in A549 cells. d) Quantitative analysis of intracellular GSH levels in A549 cells after dynamic treatments, based on western blotting shown in Supplementary Fig. 10. e) Fluorescent micrographs of Ki67 protein expression in A549 cells after dynamic treatments. Nucleus was visualized with Hoechst 33342 (10 µg mL^-1^). (M, Merge; H, Hoechst; Ki, Ki67) Scale bars represent 20 μm. g) Intracellular levels of Ki67 protein in A549 cells after dynamic treatments. *ns*: no significance, *\*P* < 0.005, *\*\*P* < 0.01, *\*\*\*P* < 0.001 and *\*\*\*\*p* < 0.0001. Data are presented as the mean ± SD.

Remarkably, the incorporation of OXA elevated intracellular ROS levels through a cascade of reactions enhancing H_2_O_2_ production. The combined treatment with FC and OXA, both in solution (FC+OXA Sol) and within the SANTA FE OXA, led to a pronounced increase in ROS, an evidenced by the intense fluorescence. Notably, the SANTA FE OXA induced even higher ROS levels than the soluble combination. This effect was markedly diminished by an ROS scavenger (VC) and an iron chelator (deferoxamine mesylate, DFO), which resulted in decreased fluorescence, thereby the pivotal role of FC in ROS generation.

Given that ROS is implicated in oxidative damage to biological molecules, including lipids, we observed through CLSM that A549 cells treated with the SANTA FE OXA or FC+OXA Sol displayed heightened BODIPY fluorescence indicative of LPO, as opposed to the minimal signal from cells treated with PBS, FC, or OXA alone (Figure 4a, d). The SANTA FE OXA exhibited the most significant LPO response, which was attenuated by DFO treatment, confirming the involvement of lipid oxidation in the observed cellular effects.

The mechanisms contributing to these observations can be summarized as follows: 1) SANTA FE OXA efficiently internalizes into cells via CD44-targeting endocytosis. 2) The release of OXA from the OXA (IV) prodrug leads to GSH depletion and disruption of redox balance. 3) The internalized OXA triggers a chain of cascade reactions resulting in H_2_O_2_ production. 4) The release of Fe^2+^ from the FC reacts with H_2_O_2_, generating substantial amounts of ROS and instigating lipid peroxidation, thereby leading to elevated LPO levels.

GSH depletion served as an additional indicator of ferroptosis.^[57]^ In the context of tumor biochemistry, GSH acts as an intracellular bastion against free radicals, neutralizing excessive ROS and mitigating oxidative damage. In our study, we quantified GSH levels in A549 cells following dynamic administration of various formulations. The GSH level in cells treated with the FC+OXA Sol ware diminished compared with those treated with the OXA alone (Figure 4e). This suggests that the FC catalyzed ROS generation via the Fenton reaction, leading to GSH depletion. Moreover, the enhanced cellular uptake of FC and subsequent release of the OXA (IV) prodrug within the micelle group resulted a more significant GSH depletion relative to the FC+OXA Sol group. These results suggest that the efficiency of SANTA FE OXA-induced ROS in depleting intracellular GSH levels.

GPX4, a pivotal regulator of lipid peroxidation within the ferroptosis pathway, plays significant bearing on the therapeutic efficacy of ferroptotic agents. Therefore, downregulation of GPX4 is synonymous with promoting ferroptosis. To validate the efficacy of our SANTA FE OXA on GPX4 suppression, western blot analysis was employed to measure GPX4 protein expression (Figure S12, Supporting Information). The analyses revealed no significant change in GPX4 expression in the control (PBS) or OXA-treated LLC cells, aligning with expectations. Conversely, cells treated with FC+OXA Sol displayed reduced GPX4 expression, with an even more pronounced decline following treatment with SANTA FE OXA. These results confirm the efficacy of the SANTA FE OXA in modulating the ferroptosis signaling pathway via inhibiting GPX4 expression.

Characteristics such as increased ROS and LPO, reduced GSH, and diminished GPX4 expression collectively signify ferroptotic activity.^[58]^ The SANTA FE OXA triggers ferroptosis through these pathways, culminating in the disruption of the redox balance within tumor cells—a critical aspect of maintaining cellular homeostasis. Consequently, this disruption amplifies the cytotoxicity of OXA (Figure 3a, b, c) and potentially augments the therapeutic efficacy of chemotherapeutic drugs via ferroptosis.

The ability of SANTA FE OXA to disrupt redox processes in tumor microenvironments showcases their potential as a nanomedicine in cancer therapy predicated on ferroptosis. Using ToC technology, we evaluated the drug efficacy of the SANTA FE OXA, by measuring proliferation of A549 cells within the chip using Ki67 immunofluorescence and Ki67 ELISA kit (Figure 4e, f). These results indicated that while OXA aline moderately inhibited cell proliferation, the FC+OXA significantly stymied cellular proliferation. The SANTA FE OXA, with excellent cellular uptake and intracellular drug release, instigated a self-amplified cascade of ROS generation and exhibited a profound synergistic anti-cancer effect, culminating in marked reduction in cell proliferation. These results highlight the promise of integrating cascade chemotherapy and self-sensitized ferroptosis in improving tumor treatment efficacy, as evidenced by the significant drug performance improvements afforded by our nano drug delivery system.

### 3.6 In vivo anti-tumor efficacy on an LLC subcutaneous model

Motivated by the promising *in vitro* anti-cancer efficacy of SANTA FE OXA, we extended our investigation to evaluate its anti-tumor activity in LLC tumor-bearing mice (Figure 5a). Analyses of tumors (Figure 5b, c, d) revealed a significant variance among the treatment groups. The concurrent administration of FC and OXA in the FC+OXA Sol exhibited enhanced anti-tumor activity compared to OXA Sol alone, suggesting a synergistic effect between FC and OXA. Remarkably, HÀ-FC-OXA-LA Micelle displayed superior therapeutic efficacy, with potent inhibitory effects on tumor progression and avoided body weight loss in comparison with both OXA Sol and FC+OXA Sol groups. This therapeutic superiority is likely due to the multifunctional attributes, encompassing robust self-assembly, excellent stability, efficient cellular uptake, and remarkable tumor targeting. Furthermore, this design of micelle facilitates site-specific drug release via reductive and enzymatic activation, amplifying the anti-tumor potency. Additionally, it possesses a self-amplified ability of ROS generation, achieving comprehensive anti-tumor effects through combined cascade chemotherapy and self-sensitized ferroptosis.

**Figure 5.**
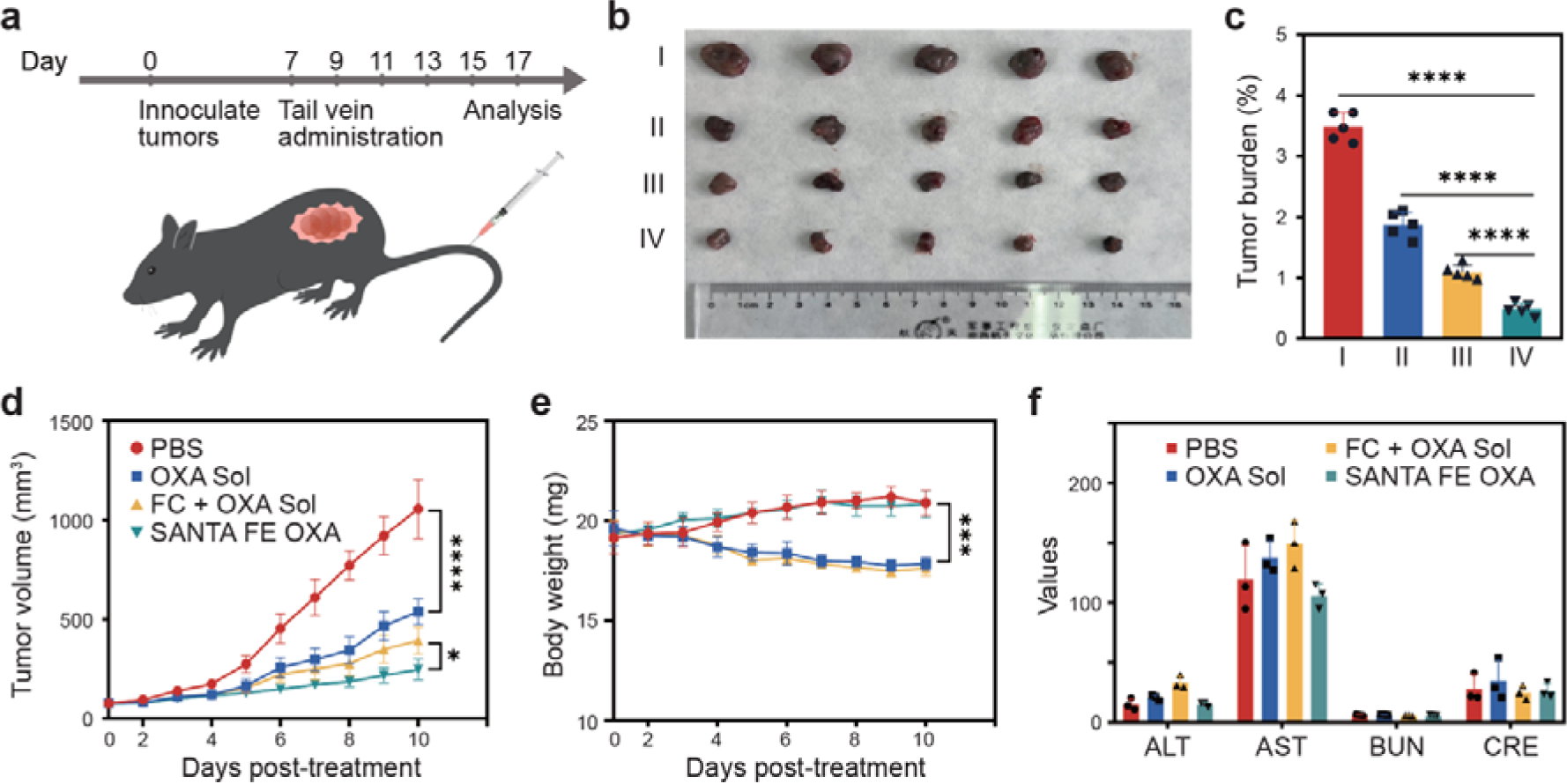
Evaluation of *in vivo* anti-tumor efficacy with LLC subcutaneous model. a) Schematic diagram of the therapeutic process of the LLC subcutaneous model, and mice were intravenously injected with 4 groups (I: PBS, II: OXA Sol, III: FC+OXA Sol, IV: SANTA FE OXA) for 5 times (*n* = 5). b) A photograph of the tumors resected from the mice after treatments. c) A bar graph of tumor burden of □-□ groups after treatments. d) Tumor volume changes of □-□ groups after treatments. e) Body weights changes of the mice in □-□ groups after treatments. f) Hepatic and renal function indicators, aminotransferase (ALT), aspartate aminotransferase (AST), urea nitrogen (BUN) and creatinine (CREA), of mice-bearing tumor after the last treatment (*n* = 3). *ns*: no significance, *\*P* < 0.005, *\*\*P* < 0.01, *\*\*\*P* < 0.001 and *\*\*\*\*p* < 0.0001. Data are presented as the mean ± SD.

Finally, the therapeutic safety profile of SANTA FE OXA was also explored. We assessed systemic toxicity by monitoring the body weight changes over the course of treatment. As shown in Figure 5e, the administration of OXA Sol and FC+OXA Sol resulted in a serious weight loss because of the nonspecific systematic distribution of the free drugs. In contrast, SANTA FE OXA had had a negligible impact on the body weight of the LLC tumor-bearing mice, at equivalent FC and OXA dosages, underscoring its enhanced biocompatibility. To broaden our safety evaluation, blood samples were collected on Day 10 post-intravenous administration for blood biochemistry and hematology analyses (Figure 5f). The results indicated no obvious alternations in liver and kidney functions, suggesting that SANTA FE OXA does not impart overt systemic toxicity under the tested conditions.

While our findings offer considerable promise, they are not without their challenges that warrant further investigation. Despite the innovative use of oxaliplatin (OXA) prodrugs within SANTA FE OXA designed for activation within the tumor microenvironment, OXA is not exempt from the typical limitations associated with many chemotherapeutic agents. Notably, OXA can induce significant adverse effects and is prone to drug resistance,^[59]^ issues that are currently the focus of extensive research.^[60]^ In our study, we successfully mitigated the side effects to a degree; however, the problem of drug resistance remains unresolved. Glutathione (GSH) is a key factor in the development of resistance to OXA-based treatments. Although our SANTA FE OXA effectively depleted intracellular GSH to induce self-sensitized ferroptosis, extracellularly secreted GSH may still facilitate resistance. This necessitates additional long-term evaluations to monitor for potential tumor recurrence and to assess the enduring efficacy of the treatment.

This study introduces a SANTA FE OXA for tumor cascade chemotherapy and self-sensitizing ferroptosis. However, the utilization of the polymer coupling method in obtaining SANTA FE OXA limited accurate control over the ratio of FC and OXA, resulting in only a fixed ratio being adopted. The investigation into the optimal ratio for achieving a synergistic anti-tumor effect between these two compounds was not systematically conducted, suggesting that further research is necessary in this regard. Additionally, since the evaluation of uptake, ROS, LPO, etc., was conducted using chips with limited cell capacity, only semi-quantitative methods were employed for characterization purposes. More precise quantitative characterization techniques could provide a more accurate foundation for this paper. Furthermore, while promising results were demonstrated by the SANTA FE OXA in 2D cells as well as in 3D dynamic tumor chips and ectopic tumor models in vivo; But if utilizing more complex orthotopic tumor models might offer stronger support for this study.

## 4. Conclusions

In conclusion, we have successfully developed a self-assembled SANTA FE OXA with potent anti-tumor capabilities, utilizing a dual-action therapeutic strategy that integrates cascade chemotherapy with ferroptosis. This SANTA FE OXA is engineered by conjugating the traditional chemotherapy drug OXA and iron catalyst FC onto HA, with integration of EG-LA. Within the tumor microenvironments, the presence of HAase and elevated GSH levels trigger the swift disintegration of the SANTA FE OXA to release FC and OXA. The released FC triggers the Fenton reaction, leading to increased LPOs and the induction of ferroptosis. Concurrently, OXA inflicts DNA damage and promote H_2_O_2_ generation via a cascade reaction, enhancing anti-tumor efficacy. Furthermore, GSH depletion exacerbates LPO levels, potentiating ferroptosis by inhibiting GPX4 expression. This innovative approach to ferroptosis induction has shown substantial efficacy in both 2D cell culture and more sophisticated 3D dynamic tumor models, as well as *in vivo* studies. Overall, this combined strategy offers a promising approach to enhance the effectiveness of chemotherapy via ferroptosis for advancing tumor suppression.

## Data availability

All relevant data are available from the authors.

## Declaration of competing interest

The authors declare that they have no known competing financial interests or personal relationships that could have appeared to influence the work reported in this paper.

## Acknowledgements

J.S. and W.M. contributed equally to this work. Funding was generously provided by the National Natural Science Foundation of China (82104109), Liaoning Provincial Department of Education Program (LJKMZ20221353 and LJKZ0940), Natural Science Foundation of Liaoning Province (2022-BS-158) and the Japan Society for the Promotion of Science (JSPS; 21H01728). The WPI-iCeMS is supported by the World Premier International Research Centre Initiative (WPI), MEXT, Japan.

## Author contributions

Conceptualization: J.S., W.M., C.T., K.K.

Methodology: J.S., W.M., S.Z., F.X., X.L., A.K., C.T., K.K.

Investigation: J.S., W.M., C.T., K.K.

Visualization: J.S., W.M.

Funding acquisition: C.T., K.K.

Project administration: C.T., K.K.

Supervision: C.T., K.K.

Writing – original draft: J.S., W.M., C.T., K.K.

Writing – review & editing: J.S., W.M., C.T., K.K.

## Notes

### Competing Interest Statement

The authors have declared no competing interest.

