## Supplementary Information for "SANTA FE OXA: Self-assembled oxaliplatin nanomicelle for enhanced cascade cancer chemotherapy via self-sensitized ferroptosis"

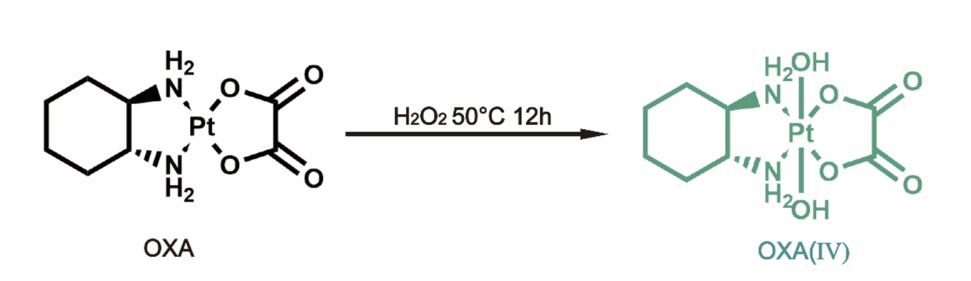


**Figure S1.** Synthesis step and structures of OXA(IV)

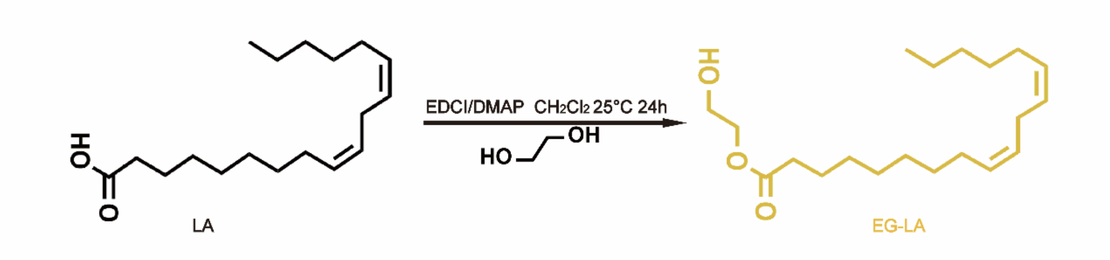


**Figure S2.** Synthesis step and structures of EG-LA

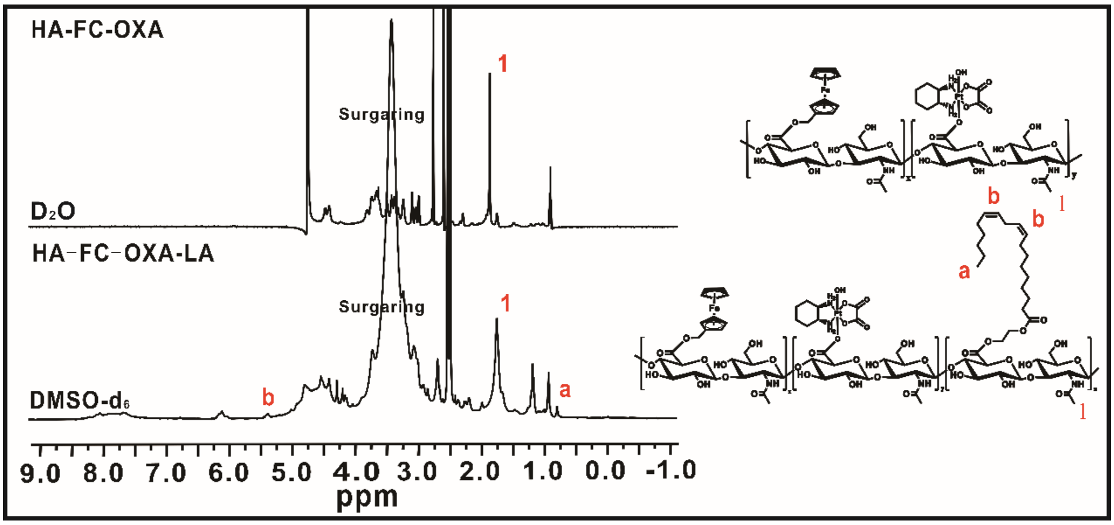


**Figure S3.**[1]H NMR spectrum of HA-FC-OXA-LA conjugates. Chemical shifts 1 (at 1.9 ppm) and a (at 0.85 ppm) represents N-acetyl group of HA and LA, respectively.

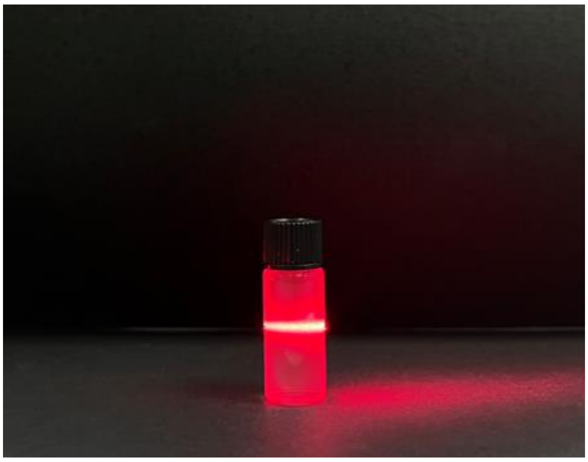

**Figure S4.** Tyndall effect of SANTA FE OXA (6mg $mL^{-1}$).

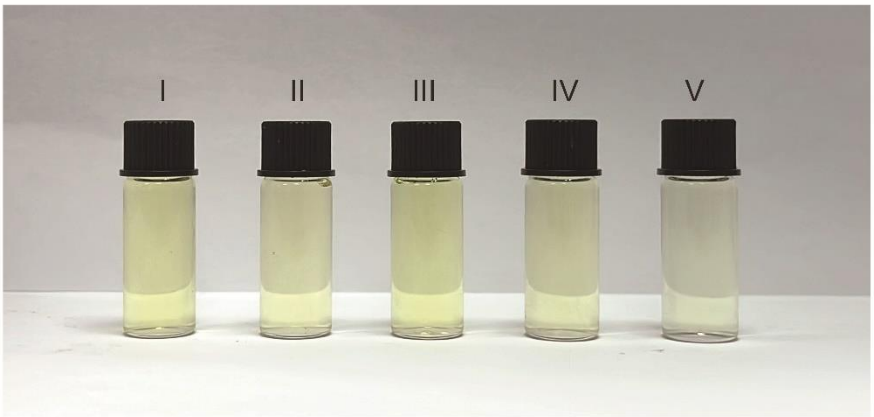


**Figure S5.** Images of DTNB solution post the reaction between various formulations (I, PBS; II, FC Sol; III, OXA Sol; III, FC+OXA Sol; V, SANTA FE OXA) and GSH. The color change reflected the level of GSH (yellow). GSH detection at 420 nm.

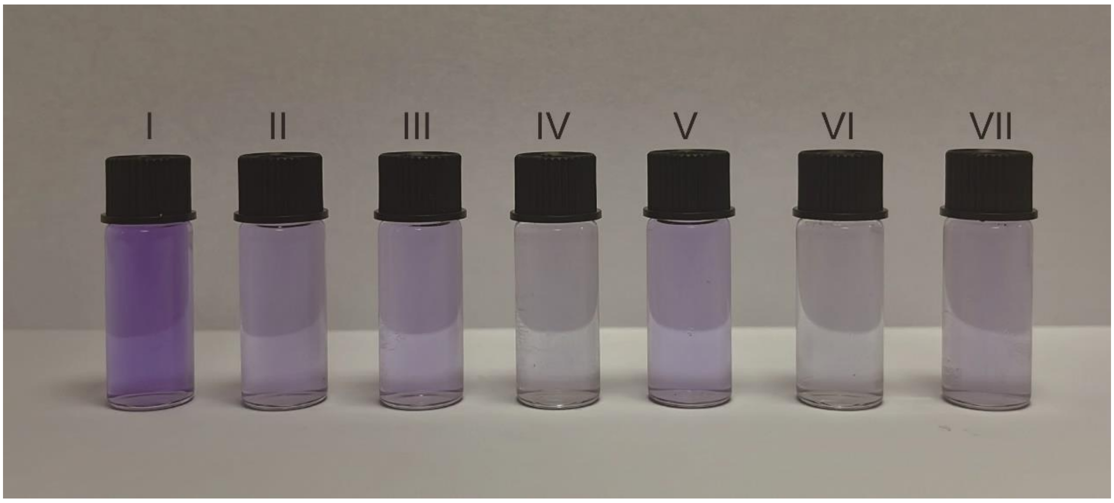


**Figure S6.** Images of crystal violet solution post the reaction between various formulations (I, PBS; II, $H_2O_2$; III, Methanol+$H_2O_2$; III, OXA+ $H_2O_2$; V, FC+ $H_2O_2$;VI, FC+OXA+ $H_2O_2$; VII, SANTA FE OXA+ $H_2O_2$) and $H_2O_2$. The color change reflected the level of •OH (violet). •OH detection at 580 nm.

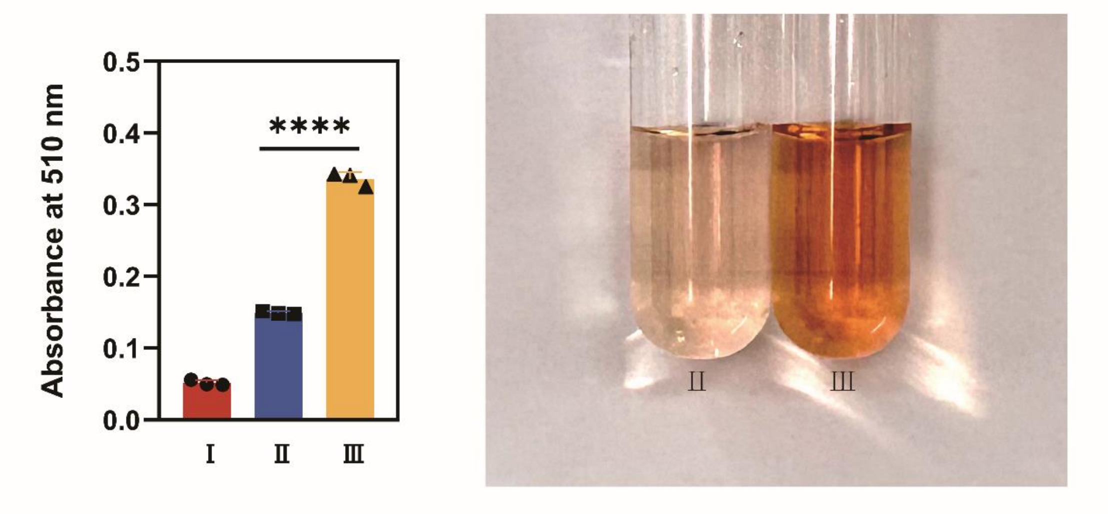


**Figure S7.** A Bar graph of $Fe^{2+}$ detection at 510 nm (*Left*). Images of $[Fe(phenanthroline)^{3}]^{2+}$ solution post the reaction between various groups (I, PBS; II, SANTA FE OXA; III, SANTA FE OXA+$H_2O_2$) (*Right*). The color change reflected the level of $Fe^{2+}$.

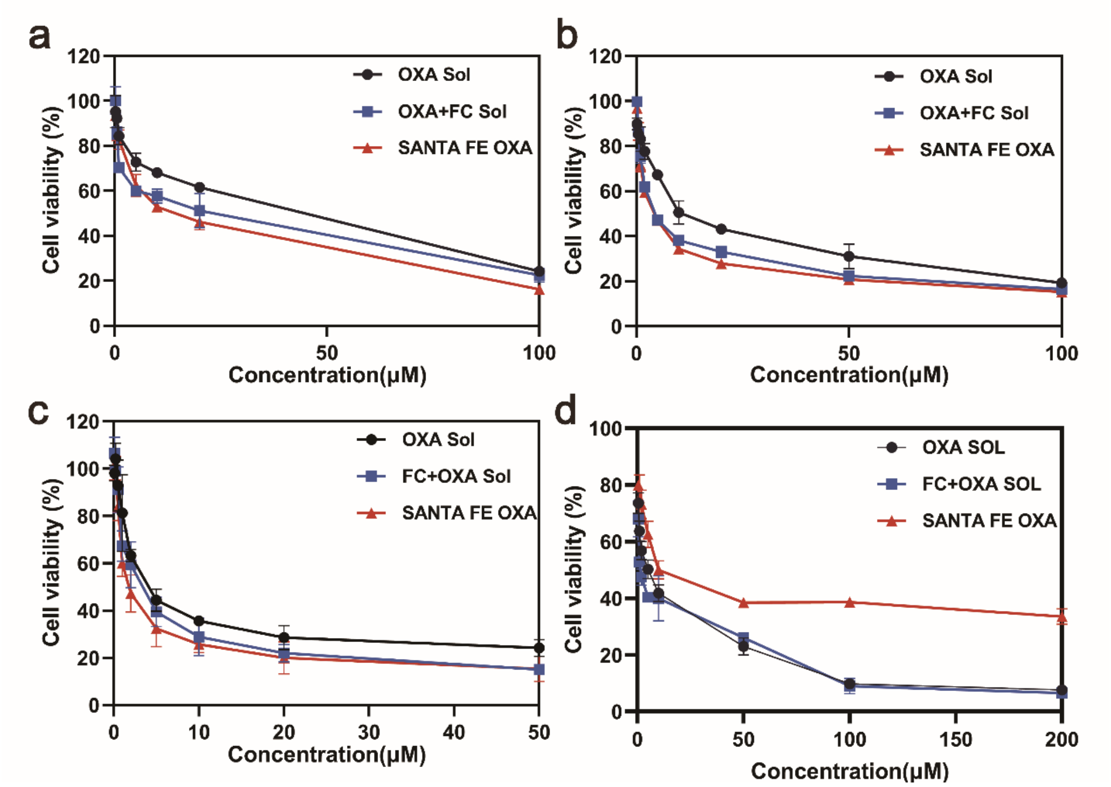


**Figure S8.** (a, b, c, d) In vitro cytotoxicity against A549, LLC, MDA-MB-231 and NIH3T3 cells treated with various formulations.

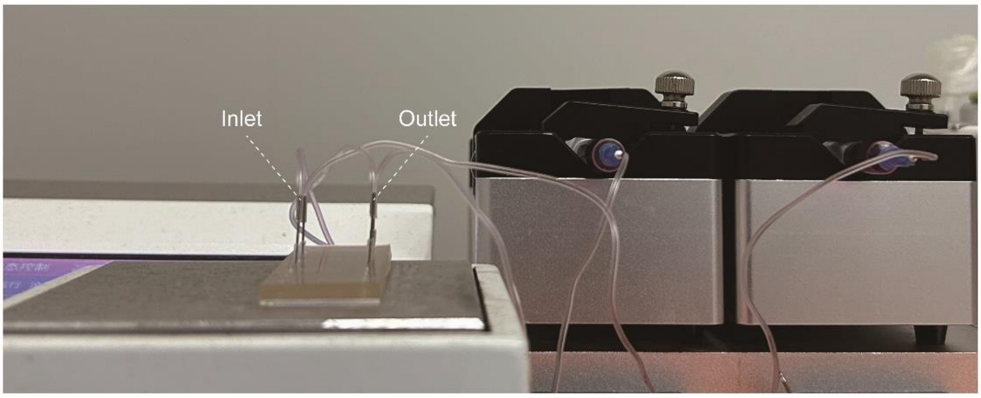


**Figure S9.** Image of the dynamic flow perfusion system. The left exit of channels was connected with a syringe pump that generates a bidirectional flow, while the right exit of channels was connected to reservoirs

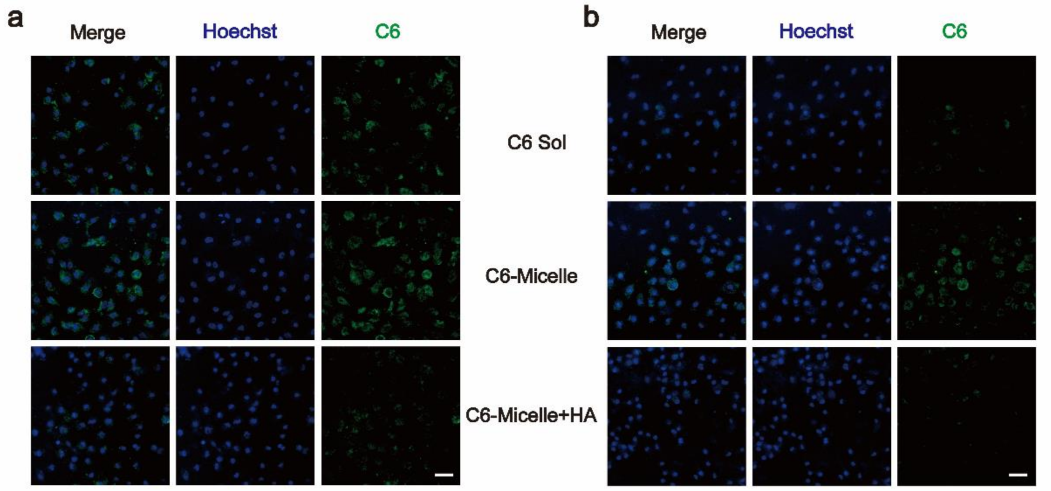


**Figure S10.** a) Fluorescent micrographs of A549 cells under on-chip static cellular uptake at 1 h. Scale bar represents 20 μm. b) Fluorescent micrographs of A549 cells under on-chip dynamic cellular uptake in A549 cells at 1 h. Scale bar represents 20 μm.

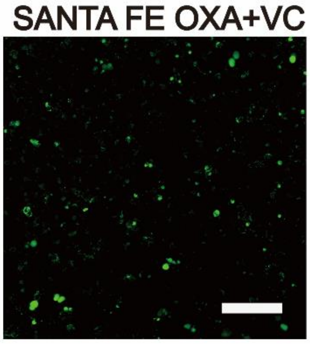


**Figure S11.** Intracellular levels of ROS in A549 cells after dynamic treatments, visualized by DCFH-DA. Scale bars represent 100 μm.

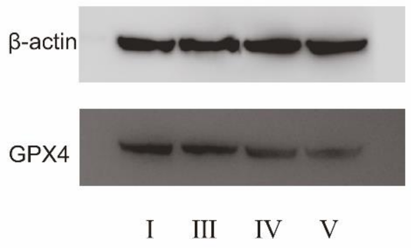


**Figure S12.** Western blotting analysis results of intracellular GPX4 in in LLC cells after static treatments with various formulations. (I, PBS; III, OXA Sol; III, FC+OXA Sol; V, SANTA FE OXA)
